# A novel method accounting for predictor uncertainty and model transferability of invasive species distribution models

**DOI:** 10.1101/2022.03.14.483865

**Authors:** Matt Hill, Peter Caley, James Camac, Jane Elith, Simon Barry

## Abstract

Predicting novel ranges of non-native species is a critical component to understanding the biosecurity threat posed by pests and diseases on economic, environmental and social assets. Species distribution models (SDMs) are often employed to predict the potential ranges of exotic pests and diseases in novel environments and geographic space. To date, researchers have focused on model complexity, data available for model fitting, the size of the geographic area to be considered and how the choice of model impacts results. These investigations are coupled with considerable examination of how model evaluation methods and test scores are influenced by these choices. An area that remains under-discussed is how to account for uncertainty in predictor selection while also selecting variables that increase a model’s ability to predict to novel environments (model transferability).

Here we propose a novel method to finesse this problem by using multiple simple (bivariate) models to search for the candidate sets of predictor variables that are likely to produce transferable models. Once identified, each set is then used to construct 2-dimensional niche envelopes of pest presence/absence. This process ultimately results in a number of possible models that can be used to predict pest potential distributions, however, rather than relying on a single model, we ensemble these models in an attempt to account for predictor uncertainty. We apply this method to both virtual species and real species data, and find that it generally performs well against conventional approaches for statistically fitting numerous variables in a single model. While our methods only consider simple ecological relationships of species to environmental predictors, they allow for increased model transferability because they reduce the likelihood of over-fitting and collinearity issues. Simple models are also likely to be more conservative (over-predict potential distributions) relative to complex models containing many covariates – making them more appropriate for risk-averse applications such as biosecurity. The approach we have explored transforms a model selection problem, for which there is no true correct answer amongst the typically distal covariates on offer, to one of model uncertainty. We argue that increased model transferability at the expense of model interpretation is perhaps more important for effective rapid predictions and management of non-native species and biological invasions.

## Introduction

Evaluating the likelihood and potential extent of exotic pest and pathogen establishment requires an accurate prediction of their potential distribution (Sutherst, 2014). A typical avenue for predicting the novel ranges of non-native or range-expanding species is to use correlative species distribution models (SDMs) (Elith et al., 2010; Jiménez-Valverde et al., 2011; Booth et al., 2014). These models attempt to estimate where a species may persist in a landscape by correlating known occurrences against a range of environmental covariates (or predictors). A vast array of SDM techniques are available, and a range of opinions on how to apply them, making defining best practice across the variety of model applications challenging (Brodie et al., 2019; Pearson and Dawson, 2003; Merow et al., 2013).

While SDM methods are particularly useful for many applications, model transferability is one area where they are known to perform unreliably. Transferability describes the ability of a model built for one situation (i.e. for a given species, in a given area over a given time span) to predict to another (e.g. under future conditions or to new environments) (Sequeira et al., 2018; Yates et al., 2018). For pest and pathogen species this may be thought of as extrapolating a model fitted to data from the native range (or another established invasive range) to a novel area that may be at risk of that species establishing. Inherently, transferability is critical to making accurate biosecurity decisions because the ability of models to predict under new scenarios forms the baseline of many risk assessments and for informing tactical decisions such as where to allocate finite surveillance resources (Guillera-Arroita et al., 2015; Vall-llosera et al., 2017) There are many components that may affect a model’s ability to transfer well (as discussed in detail in Sequeira et al. (2018); Table 1), but one that is often not dealt with is choice of predictor variables (but see Synes and Osborne (2011)).

**Table 1:** List of species used in this study. ‘Native’ are the number of points from the native range, ‘Invasive’ are the numbers of points from the non-native range(s). ‘Expansion’ and ‘Unfilling’ refer to the niche change metrics reported in Hill et al. (2017), and whether this process is evident for a given species. Briefly, expansion refers to the species exhibiting niche expansion in the non-native ranges, whereas unfilling referes to the species occupying environmental space not yet realised in the non-native ranges.

| Order | Species | Native | Invasive | Expansion | Unfilling |
| --- | --- | --- | --- | --- | --- |
| Acari | *Halotydeus destructor* | 42 | 463 | + | + |
| Coleoptera | *Anoplophora glabripennis* | 28 | 55 |  | + |
|  | *Atrichonotus taeniatulus* | 31 | 22 | + | + |
|  | *Aramigus tessellatus* | 69 | 42 | + | + |
|  | *Leptinotarsa decemlineata* | 164 | 246 |  | + |
|  | *Harmonia axyridis* | 109 | 1340 | + | + |
| Diptera | *Aedes albopictus* | 396 | 2441 | + |  |
|  | *Bactrocera dorsalis* | 186 | 386 |  | + |
|  | *Ceratitis capitata* | 273 | 353 |  | + |
|  | *Tipula oleracea* | 54 | 251 | + | + |
|  | *Tipula paludosa* | 55 | 266 |  | + |
|  | *Zaprionus indianus* | 65 | 318 |  |  |
| Hemiptera | *Phenacoccus manihoti* | 144 | 31 |  | + |
|  | *Thaumastocoris peregrinus* | 43 | 214 | + | + |
| Hymenoptera | *Linepithema humile* | 123 | 1020 | + |  |
|  | *Pachycondyla chinensis* | 39 | 71 | + | + |
|  | *Pheidole megacephala* | 24 | 165 | + |  |
|  | *Solenopsis invicta* | 184 | 1099 | + | + |
|  | *Tetramorium* species E | 40 | 52 | + |  |
|  | *Tetramorium tsushimae* | 58 | 25 |  | + |
|  | *Wasmannia auropunctata* | 215 | 117 |  |  |
| Lepidoptera | *Epiphyas postvittana* | 203 | 35 |  |  |

Causal variables are critical to accurately predicting the distribution of a species to novel environments, such as for non-native species into new environments or under climate change scenarios (Varela et al., 2011; Austin and Van Niel, 2011; Fourcade et al., 2017). If the predictor variables are indirectly associated with defining a species’ niche, then the model will only be able to predict the populations in which it was trained on (Austin, 2002) and unreliably predict elsewhere. (Randin et al., 2006). This is because correlation structures between non-causal variables may change between geographic locations (Dormann et al., 2013), and among different populations. This in turn, can diminish a model’s ability to predict to species persistence in new regions and environments. However, as it is tremendously difficult to ‘know’ all the causal variables governing a species distribution, determining which predictors should be included in a model and how best to justify them is a significant challenge (Barbet-Massin and Jetz, 2014).

A range of studies have attempted to determine which predictor variables are best to include in a model, and in lieu of truly causal predictors, how proximal these variables may be (e.g. Petitpierre et al., 2017; Syphard and Franklin, 2009; Barry et al., 2015). Despite this, there has been little attempt to review or develop methods to simply identify “good predictive variables” from the suite typically available to modellers and how to best use them to reduce errors that may be associated with model transferability.

Typically, the data used in SDMs are based on *ad hoc* observations of species presence. Inherent with these data are often strong sampling biases that can be caused by a number of factors (e.g. remoteness, survey resources, public interest) that may result in poor or patchy species records across geographic and environmental gradients. By fitting complex models to such data, there is a significant risk the model will fit these sampling biases as opposed to what we are interested in – the relationships between predictor variables and species persistence (Merow et al., 2014). This is a strong argument for focusing on simple models. Envelope based methods provide the simplest form of SDMs, with simple functions (e.g. bounds and step features) defining their response variables (Elith et al., 2005; Merow et al., 2014), and while they may not be as ecologically interpretable as other methods that estimate response curves (e.g. regression methods), they should be less prone to overfitting.

To address these issues of selecting predictor variables and increasing model transferability, it may be practical to select sets of variables based on how useful they are at defining the species known distribution, and then represent uncertainty across these putative proximal variable sets. In a recent paper, Breiner et al. (2015) used “ensembles of small models” to predict the distributions of rare species. Breiner et al. (2015) found that ensembling an array of simple models, each with only two variables, outperformed standard SDM methods that had often had more complex sets of predictors. While these models were not tested for transferability into novel environments, it demonstrated that the use of many small models was able to outperform fitting a single, more complex model. It follows then, that by “mining” for the most useful sets of variables using simple (i.e. bivariate) models it should be possible to identify which sets of predictors perform better and then ensemble the *best* sets into a single map that conveys the uncertainty in predictor set selection. This approach may avoid some of the issues associated with transferability, multi-dimensionality and correlation issues that “complex” SDMs often exhibit.

Here we examine the ability of an approach that incorporates an ensemble of simple models and envelopes to account for predictor uncertainty. Our approach ultimately recasts the process of constructing a useful SDM as a predictor selection uncertainty problem, as opposed to one of model selection (with the implicit paradigm that there exists a “best” model). We first test this approach using virtual species where we know the environmental limits and therefore can assess model transferability against truth. We then apply this method to a number of insect species that have successfully invaded new geographic regions.

## Methods

### Environmental predictors

There are a number of different environmental predictors that are at the appropriate grain and extent needed to fit SDMs, and many of these are based around climate. While different landscape and anthropogenic variables can also be included, climatic variables are most easily accessible and generally considered a particularly good set of indicators of species distributions at a broad geographic scale, especially when little is known about the species biology and ecology. Employing climate in this way assumes that some of the climate variables are proximal to processes defining the species distribution. Several databases provide global long-term climatic gridded datasets of precipitation and maximum and minimum temperatures, appropriate for species distribution modelling at large geographic scales. For this study, we used the 19 bioclimatic variables from WorldClim 2.0 (Fick and Hijmans, 2017) at a 10’ resolution. This set of 19 variables was developed for use with the BIOCLIM package in 1996 (see, for example, Lindenmayer et al. (1996)), which also pioneered the thin-plate spline interpolation methods used to build the WorldClim data. From here on we call these data the (lowercase) “bioclim” set, as they are derived as part of the BIOCLIM (Nix, 1986) package. BIOCLIM (from here on (uppercase) “BIOCLIM” refers to the modelling method) was instrumental in not only defining the role of SDMs, but also in the description of these variables that are used widely today (Booth et al., 2014). These data provide an easily accessible, and therefore commonly employed modelling resource.

### Model construction

Here our goal was to build models from native range occurrence data, and then use these models to predict to the invasive (or non-native) range. While there are good arguments for fitting models using both native and invasive range data to predict potential distributions, here we are concerned about the ability of different modelling methods to predict the potential invasive range from native occurrence data alone. To investigate transferability across a range of approaches, we constructed SDMs using convex hulls, bounding boxes, range bagging and the commonly used machine learning tool, Maxent. These methods were employed under two broad categories: ensembles of simple (e.g. Convex Hulls, Bounding Boxes, Range Bagging, Maxent) models, and single multivariate models (Maxent). We applied all methods to two sets of data, first virtual species data (simulated using the methods described later) and second, actual species data that has been used previously in the investigation of SDM performance for invasive species.

#### Predictor selection

To construct our ensembles, we chose simple “envelope” methods that classify environments (predictor variable ranges) as either suitable (inside the envelope) or unsuitable (outside the envelope). Specifically, we constructed many two variable envelopes of species presence within their native range using pairwise combinations from the bioclim variable set. Instead of using all combinations of variables, we restricted the combinations to one temperature (bioclim 1-11) and one precipitation (bioclim 12-19) variable. The rationale here, is that at least one aspect of temperature and one aspect of precipitation is likely to be limiting a species potential distribution (Figure 1).

**Figure 1:**
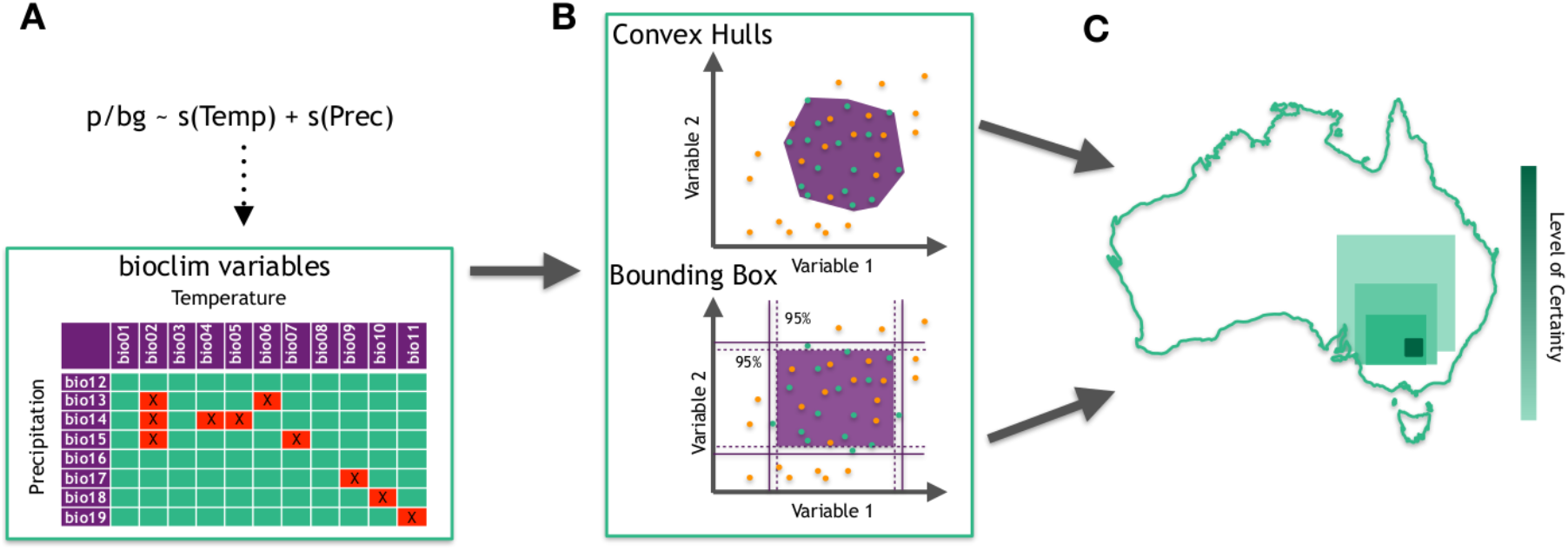
Schematic of new ensemble modelling technique. **A**. Variable pairs of temperature and precipitation using the bioclim variable are used in simple GAMs and then assessed for relative model fit using AUC (and other metrics, see text). Green squares are the pairs kept, red squares with crosses are omitted. **B**. The variable pairs selected are used to create two-dimensional environmental envelopes capturing all the presence points (green points) inside all available environments (green + orange points), using a minimum convex polygon (convex hulls (CHE)), or by use of bounding boxes defined through the BIOCLIM algorithm (BBE). **C**. Each of the two-variable envelopes from B are projected to a new geographical surface (as a gridded raster) and then stacked (summed. The results are then averaged, and the resulting surface is a continuous scale from 0-1 reflecting the level of certainty (closer to 1) that a given raster cell falls in the environmental limits across the given predictor variables.

In order to identify which set of variable pairs were most closely strongly related to the species’ native distribution, we fitted bivariate Generalised Additive Models (GAMs) with a Point Process Model (PPM) implementation for presence-only data (Renner and Warton, 2013). GAMs were constructed as a down-weighted Poisson regression (DWPR) and smooth terms were restricted to the 2^nd^ order polynomial at the most, ensuring simple curves are fitted for responses – and thus reducing possible over fitting. Terms were either fitted additively or jointly. Joint fits allow interactions between predictors, and can be achieved in the `mgcv` package in `R` by fitting a smooth surface in two dimensions over the chosen variables rather than two one-dimensional fits, one for each variable. We built models using presence-background data, which are typically the most readily available data for non-native species. Background points provide a sample of the environment in the region from which presence points are available, here they were all the cells across the geographic extent of the calibration (or native) range.

Relative performance of GAMs were estimated for all pairs of variable combinations. That is to say, we compared the performance of variable pairs only against one another, and not on the absolute predictive performance of the model (which is akin to model selection). To determine performance of variable pairs, we calculated AUC for each variable pair, and then discounted the “worst” performing models by discarding the bottom quantile (25%) of variable pairs. Using the retained variable pairs (i.e. top 75% quantile), we then used two envelope methods (Convex Hulls and Bounding boxes) to classify the environmental space within each predictor pair.

#### CHE: Convex Hull ensembles

We created convex hulls around the presence points (of the training, native range data) in environmental space, and then projected these envelopes onto the geographic space of the native range, and the invasive (non-native) range. Each envelope gives binary values of 1 for inside the hull, and 0 for outside of the hull. The resulting geographic layers for each model pair were combined together and then averaged to produce a single, continuous, prediction between 0 and 1 for the CHE. Here 0 means that a grid cell is outside all of the available pair-based envelopes (not suitable), and 1 means that the grid cell has environmental space inside all of the pair-based envelopes (highly suitable).

#### BBE: bounding box ensembles

To further examine the ensemble approach we used bounding boxes (BBE: bounding box ensembles), drawing from one of the early SDM methods, BIOCLIM (Nix, 1986). BIOCLIM works by creating a bounding box in multidimensional environmental space, and defines core and marginal environments for the species in relation to the predictors used by examining percentile distributions. Core environments are defined as the environments in which all points fall inside the 5 and 95 percentile range for all the predictors used (from an available set of 12 in the case of Nix (1986), later extended to 19 and then 35 (Booth et al., 2014)). Marginal environments are defined as where the values fall outside the 5 and 95 percentile range, but not the limits, for one or more of the predictors. In this way, the distribution is driven by the most limiting variable. BIOCLIM is typically used with several predictor variables, and here we started again with the 19 available from WorldClim (Fick and Hijmans, 2017). We used the implementation of BIOCLIM in the *dismo* package in R (Hijmans et al., 2015), to create multiple, two-variable models, using a pairwise approach across the bioclim variables available (one from 1-11, one from 12-19) (Figure 1). We used the AUC scores from the GAMs again to omit the bottom 25% of performing variable pairs. To compare how the BBE approach may perform in regards to a ‘default’ BIOCLIM model, we also ran a BIOCLIM model using all 19 available predictors and projected this to the non-native range.

#### Maxent

To compare how our approaches stack up against a popular SDM approach, we ran three types of Maxent models Phillips et al. (2006) using version 3.4.0. Firstly, we ran a model with the complete set of bioclim variables. Maxent has internal predictor selection mechanisms, and although we expect a complex model like this to be overfitted, it should provide a good comparison based on making no choice in predictor inclusion. Secondly, we ran Maxent models using eight predictor variables that have been used elsewhere for invertebrates (see Hill et al. (2017)), sometimes referred to as “state of the art” (Petitpierre et al., 2017). While the fit and performance of Maxent models can be improved through exploration of different features and parameters, we chose to leave all at default for two reasons: 1) to provide a baseline, and; 2) because default settings are common throughout the literature (despite issues with this choice (Merow et al., 2013)). Thirdly, we used Maxent to build simple two-variable models in the same framework as our CHE and BBE approaches. In an attempt to avoid overfitting for these models, we did not allow hinge, product or threshold features (models could only use linear or quadratic terms). The same variable restrictions (one temperature, one precipitation) and performance assessment using AUC scores based on the GAMs fitted to native range data applied, and the resulting single layer was an average across all retained Maxent models.

#### Range bagging

Range bagging was recently proposed by Drake (2015). This algorithm uses presence-only data to estimate the environmental limits of species habitat by subsetting the multidimensional environments (to user-defined levels of dimensionality), and then using convex hulls to estimate boundaries in each subset of environmental dimensions. Range bagging fits models to the individual samples and averages the outcome by using votes (how often a given environment occurs inside niche boundaries) on the ranges of convex hulls obtained from bootstrap samples across all the environmental dimensions. In this way, it is an approach that shares some similarities with our CHE and BBE methods. However an important difference here is that the range bagging method does not use the GAMs to rank variable pairs.

The approach has seen recent applications to invasion biology, and appears promising for biosecurity applications (Camac et al., 2019). Part of the appeal for this approach is that only presence data are needed, and no absences or background data are required, and thus, reduces the number of subjective decisions required in the modelling process. We fit range bagging models using code from Drake (2015), using for the presence-only data of the native range, and set range bagging to subset using only two dimensions to be consistent with our other approaches. In addition to running range bagging at default settings (all variables pairwise, and in two dimensions) using code as supplied by Drake (2015), we made one modification to the range bagging method, and required it to use variable pairs in the same way as our CHE and BBE methods (1 from the bioclim temperature variables 1-11, and 1 from the bioclim precipitation variables 12-19). We then projected our range bagging model to the non-native region for each species.

### Fitting datasets

#### Virtual species

To test how different modelling approaches performed at predicting the potential distribution of a species, and thus how transferable they were, we generated virtual species. Virtual species allows us to know the “truth” behind a distribution in both “native” and “non-native” (i.e. invaded) ranges, and allow us to compare predictions made by various SDM techniques to assess the accuracy and transferability of each method. There are a number of ways for generating virtual species distribution across the landscape (e.g. Leroy et al. (2016), but as we were interested in creating bounding boxes in environmental space between pairs of variables, we simulated species occurrences by setting upper and lower limits for each variable. The goal here was not to create ‘realistic’ species distributions, but rather geographical expressions of limits set within environmental ranges. Specifically, we focused the analysis on transferability between two distinct geographic regions, South America (native range) and Australia (non-native range). South America provides a vast range of climates that are both analogous and non-analogous to Australia, which allows us to test transferability across a range of environmental conditions and ensured that variables were not set outside of the range present across South America. We selected a random temperature variable from the bioclim set (1-11) and a random moisture variable (12-19). Virtual species distributions were created by taking the upper and lower limits of the entire surface area, and selecting random uniform distributions inside these to give upper and lower ranges for each variable. We enforced that the lower and upper bounds of the environmental range had to be taken from the respective sides of the mean, and at least one standard deviation existed between upper and lower bounds, to ensure there was at least some expression of that niche in geographic space. Within these limits (a box in the environmental space of these two variables) the species is always capable of being present. By contrast, outside of the box the species is always absent (i.e. cannot persist). Thus these virtual species exhibit pure niches, with no biotic interactions, and they have no stochastic element, i.e. the probability of presence is zero or one. The limits were set based on a random selection across the global range of environments. These “niches” were then realised in South America and Australia, which makes it possible to examine geographic ranges of environmental similarity between ranges.

For assessing how well the different modelling approaches could predict the distribution (or geographical representation of the true niche) in the non-native range (Australia), we followed the general framework but importantly, left out the two variables that were used to create the true niche. These are the causal variables and by omitting them from the analyses we are only considering the other 17 variables which have varying degrees of proximality to those. This means that we are acknowledging that we will never know the “true niche” but can only make approximations of it, based on variables that range from proximal to distal.

#### Real species data

We also examined the performance of different modelling approaches using occurrence records for the 22 invasive insect species dataset from Hill et al. (2017) (see Table 1). Many of these species are priority plant pests, disease vectors or nuisance pests. These species were chosen as they have all successfully established in new geographic locations, and provide data for both “native” and “non-native” regions to train and test models with. While these species were chosen as they have occurrences for both native and non-native regions, the underlying data are inherently biased in the way they have been sampled across different regions. These biases include under-sampling across the total range of environments, and increased sampling in areas where the species are noted pests. Another important caveat is that Hill et al. (2017) found evidence of ‘*niche shift* ‘ (adapted or expanded to novel environments), in a number of these species, suggesting that the distributions in the novel ranges may extend outside of what is predictable from training on the native range alone (Table 1). Bearing that in mind, the trends presented do encapsulate what may be expected from a dataset of actual species data, complete with inherent bias in sampling effort and geographical extent that is typical of many non-native and invasive species. We applied the same methodology for each of the species in order to see how different methods performed across a group of species of varying range size, data availability and sampling evenness.

We defined backgrounds by selecting all unique “biomes” (see Olson et al., 2001) that occurrence points intersected with, as this has proved to be a useful extent for modelling these species (Hill et al., 2017). To help limit sampling bias, the presence data were rescaled to the level of the grid cell data (10’), so that there was only one occurrence per grid cell. The background was selected within the native range as either all available cells within biomes with at least one occurrence point, or 50,000 random cells across these biomes, whichever was fewer.

### Assessing model transferability

To examine how models performed in predicting the non-native ranges, we employed three different measures. As we had independent datasets (non-native range) for each species to evaluate the models, we first employed AUC on this test dataset to assess performance. The AUC score was calculated using the non-native range distribution and the associated background (i.e. the geographical boundaries defined by the definitions of either continents, or biomes).

As different evaluation techniques may yield different results, we also used the Boyce index (Boyce et al., 2002), and sensitivity (see Petitpierre et al., 2017), as measures for the final surfaces projected into the novel ranges. The Boyce index is a presence-only evaluation technique and measures how much model predictions differ from a random distribution of observed presences across a prediction gradient (Hirzel et al., 2006). The Boyce index partitions the total habitat suitability range into a set of bins, and for each of these calculates the predicted expected ratio, using the Spearman rank correlation coefficient (Hirzel et al., 2006). The Boyce index ranges from -1 to 1, with positive values indicating good model performance, values near 0 indicating models no better than random and negative values indicating prediction of poor suitability where there are presences (Di Cola et al., 2017). Sensitivity is the percentage of true presences correctly predicted by the model.

## Results

### Virtual species

There was variation in the ability of different modelling approaches trained in South America (native) to transfer across/predict to Australia (non-native). The different applications of Maxent (using all variables, using a pairwise approach) had high AUC and Boyce index scores, whereas there was more variation in these scores for the other approaches (Fig. 2). Sensitivity was highly variable for all of the approaches, however again, the two Maxent approaches gave the highest sensitivity values, indicating that Maxent was most reliable at predicting true presences (sites inside the niche in Australia). In addition to the two Maxent approaches, the BBE approach also worked well for these virtual species, with high Boyce and good AUC (Fig. 2). Similarly, the CHE approach also worked reasonably well for the evaluation scores, but there was more variation for the Boyce scores than for BBE. The two range bagging approaches had the largest amounts of variation across the three evaluation techniques, and consequently the lowest mean scores (Fig. 2). BIOCLIM had good Boyce scores, however the AUC and Sensitivity were variable and lower than the best performing methods.

**Figure 2:**
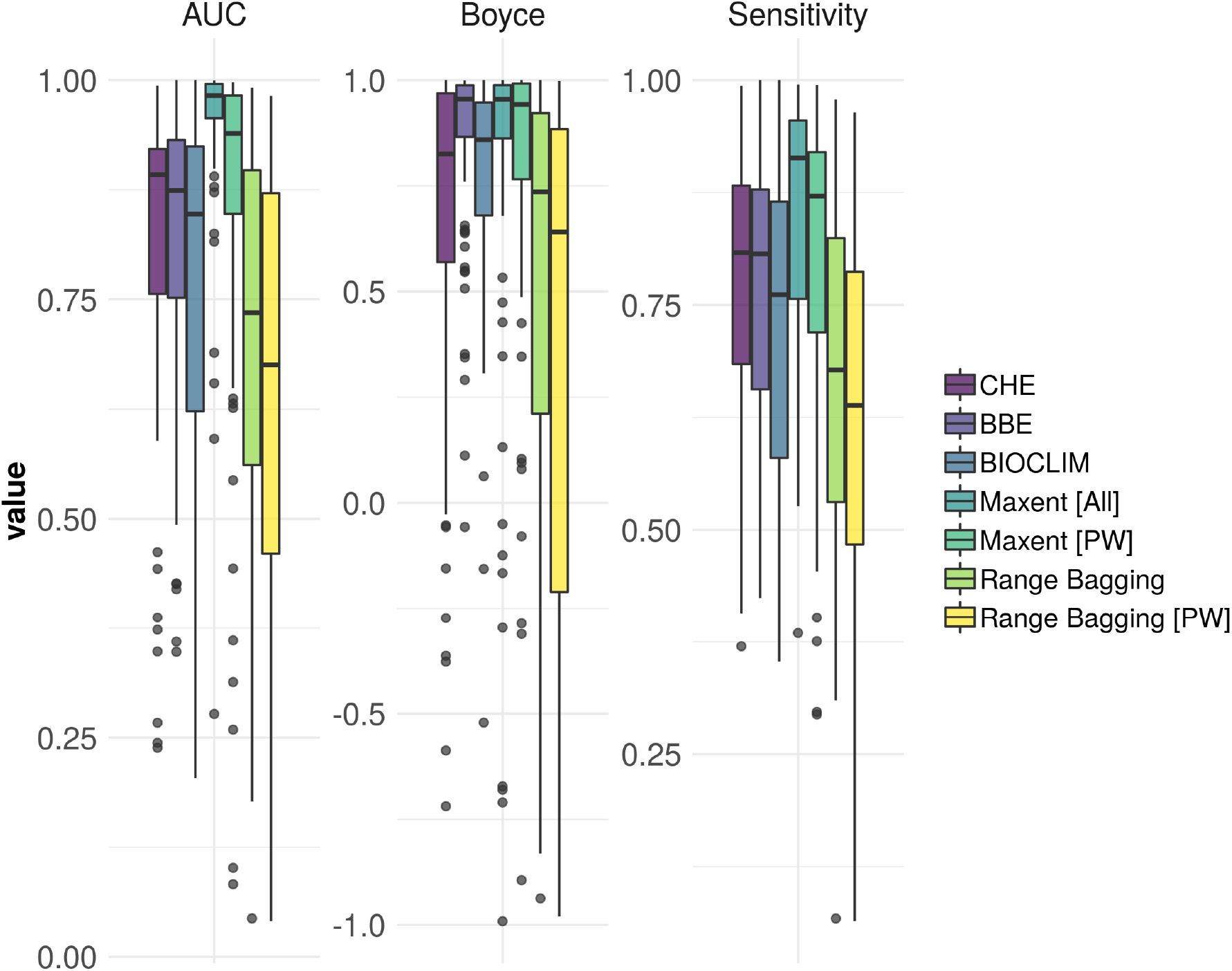
Evaluation scores for a comparison of the different SDM methods for virtual species: CHE = convex hulls ensembles; BBE = bounding box ensembles; BIOCLIM = BIOCLIM algorithm using all 19 bioclim variables (with two causal variables omitted); Range Bagging [PW] = default Range Bagging algorithm with dimensions set at two and selection of only one temperature and one precipitation variable at a time, leaving out the two causal variables; Maxent [PW] = ensemble of simple Maxent models and selection of only one temperature and one precipitation variable at a time, leaving out the two casual variables; Maxent [All] = Maxent using default settings and all 19 bioclim variables

## Real species

The real species data provides a test of projecting an incomplete understanding of the niche and the associated distributions to a novel geographic range. Here the models were calibrated on only the native range and then projected to the invasive range. Normally it would be better practise to include the invasive range in the model training (Broennimann and Guisan, 2008) — however here we are providing a test of the ability of models to project from limited data rather than a description of the entire niche.

Figure 3 shows the evaluation scores for the different SDM techniques we employed for these real species data. While there is variation within methods, across them the distribution of AUC is fairly similar (assessed on the non-native distribution and range). Likewise, Sensitivity is above 0.8 for most species and methods (except BIOCLIM) which may be considered as “being transferable” (Petitpierre et al., 2017)). The greatest variation is in the Boyce score, which places the CHE and Maxent[PW] methods as better performing than the other approaches Fig. 3). On balance across the evaluation techniques, The Maxent[PW], CHE and BBE approaches appear to work well according to all the evaluation methods employed here (Fig. 3), although most of the approaches here could are performing except for BIOCLIM and perhaps Maxent [All] which also has low Boyce scores. The range bagging approaches were similar, with the default implementation performing slightly better than the pairwise approach for AUC and sensitivity (Fig. 3).

**Figure 3:**
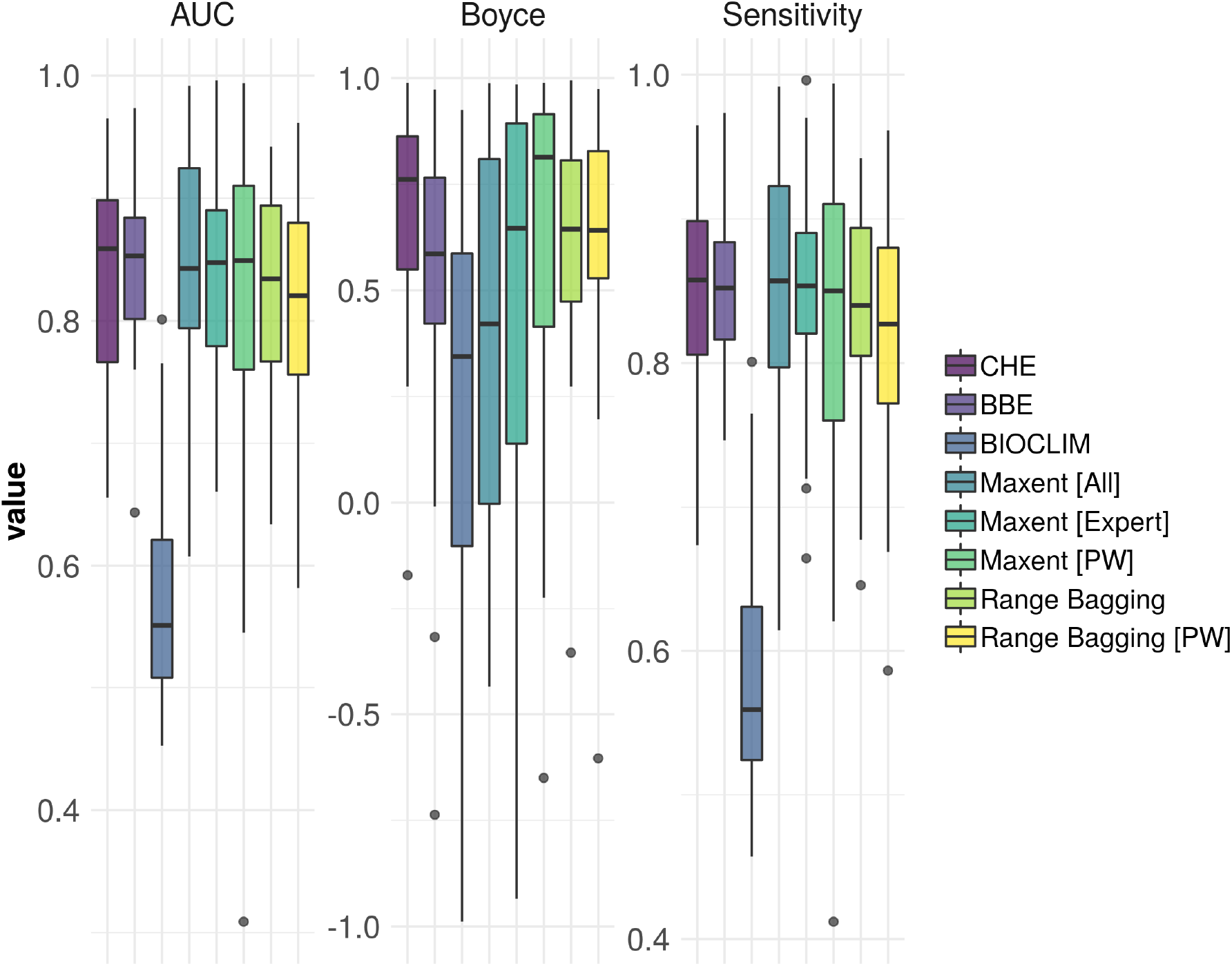
Evaluation scores for a comparison of the different SDM methods for the 22 invasive insects species data: CHE = convex hulls ensembles; BBE = bounding box ensembles; BIOCLIM = BIOCLIM algorithm using all 19 bioclim variables; Range Bagging = default Range Bagging algorithm with dimensions set at two; Range Bagging [PW] = default Range Bagging algorithm with dimensions set at two and selection of only one temperature and one precipitation variable at a time; Maxent [PW] = ensemble of simple Maxent models and selection of only one temperature and one precipitation variable at a time; Maxent [Expert] = Maxent using default settings and eight predictor variables described as useful for predicting insect distributions; Maxent [All] = Maxent using default settings and all 19 bioclim variables

## Discussion

The discussion around “best practice” for using species distribution models in pest risk mapping is typically focused on model selection, with emphasis often placed on finding the best performing models (by some statistical measure), and using that information as the basis for projection to novel areas and risk assessment (e.g. Fan et al. (2018); Jiménez-Valverde et al. (2011)). By transforming a model selection problem to one of model uncertainty across predictors, we suggest alternative approaches that perform well against established methods, and provide ease of interpretation through an uncertainty framework. Specifically, when the choice of which available predictors to use is unknown or unguided, constructing an ensemble of small (two variable) models allows for model construction to be focused on selection of “good” performing variables (ones that perform well in both native and non-native ranges), rather than relying on model fit as the indication of the “best” model. Our methods allow for putatively more proximal variables to be identified and to apply them in a simple manner to not only increase transferability (Petitpierre et al., 2017), but provide a more tractable approach to developing risk maps of potential non-native distributions.

Both CHE and the pairwise Maxent approaches were consistently good performers across virtual and real species models. Previously, using a ensemble of two-variable Maxent models has yielded useful results for predicting the distributions of rare species (Breiner et al., 2015), and here we provide evidence that this approach can be extended to a transferable modelling framework for non-native species. One important distinction here is that we used paired temperature and precipitation variables to rank and include variables in the ensemble, rather than using all available pairs across the predictor set. While ecologically it seems rational to define niche limits using pairs of temperature and precipitation variables, we also found increased variability in performance when applying this to the Maxent and range bagging approaches, an issue that may warrant further investigation. Further, we have limited our analyses to only use the bioclim variables, although other abiotic predictors may be better suited to model a given species, however these could easily be incorporated into the framework. For example, the availability of gridded global soil datasets have become more readily available and would serve as useful predictors for some taxa (see Booth (2018)). We also chose not to weight the different models or predictor pairs, as has been done in elsewhere (Breiner et al., 2015). As our goals were focused on communicating uncertainty across available predictors in an attempt to identify “good” predictors, using unweighted pairs provides a tractable and interpretable approach. As the true response to distal variables is likely to be complex (Merow et al., 2014), using an ensemble of simple models can help avoid issues associated with that complexity (Breiner et al., 2015). Overfitting can also be minimised, therefore helping to filter any environmental noise, and biased predictions associated with extrapolation (Yates et al., 2018; Bell and Schlaepfer, 2016).

There are some important differences between the CHE and pairwise Maxent approaches that are worth considering when making choices of appropriate modelling frameworks to apply to non-native species. Firstly, Maxent can be thought of as modelling closer to the realised distribution of the species, the CHE closer to the potential distribution (Jiménez-Valverde et al., 2011). This distinction is important when considering how the species-environment relationships are defined given the data available, and what this means for extrapolation. In our CHE approach the relationships are simplified to environmental limits and provide a final spatial prediction as a gridded surface that represents the frequency that a given cell fell within the environmental space as defined by the convex hulls. This is directly related to the uncertainty in predictor variables, with an increase in the number of times a grid cell falls within environmental space corresponding to an increase in certainty in the predictors retained. Alternatively, the pairwise Maxent approach models the species-environment relationships as functions in two-dimensions at a time, and then the averaged probabilities of those relationships used to predict distributions. Maxent fits the relationship to data, there is a chance that when projected to a new set of environments a combination of those variables (and their interactions) from the training data does not exist. The CHE thus provides a more conservative approach to predicting distributions, with simple environmental limits, whereas the pairwise Maxent approach allows for estimation of species’ responses to environmental predictors using probabilistic functions which may be closer to the ecological interactions - but more prone to adverse effects of extrapolation if the training data does not allow for the characterisation of that relationship.

The other approach we investigated, range bagging, also performed well for the real species data, despite having highly variable performance for the virtual species. Range bagging shares some similarities to the CHE approach, in that the environmental relationships are estimated through the use of convex hulls across the different combinations of available predictors. This means that the resulting predictions are able to be interpreted in the same manner as for CHE. While CHE and range bagging may lead to larger predictions (more conservative), this may be more preferable for invasive species (Kramer et al., 2017). Comparing modelling performance between methods is a lot more straightforward than explaining the reasons behind differences (Breiner et al., 2018), but the inconsistent performance for range bagging on the virtual species may be due to a couple of reasons. Firstly, range bagging is sub-sampling the occurrence data, so unlike the other methods here it does not utilise all available data when fitting. Secondly, and perhaps more importantly, part of the variability may come from including poor pairs of predictors, especially when the causal variable pair is absent. When looking at performance for the virtual species for all methods it is important to note that the distributions that were created are not “realistic” species distributions, and instead are focused on testing the ability to identify proximal variables in absence of the causal predictors for each given virtual species.

One of the biggest challenges of using real species datasets to evaluate modelling methods is “messy” datasets (as a result of sampling bias, under-representation of range, missing data, errors in data). For many non-native species the data available is not sampled with the same intensity and process across native and invasive ranges. Sampling processes under a real world situation may only yield few occurrence points across the actual distribution, with spatial bias in reporting. This translates into methods that may otherwise perform well, being penalised for characterising patterns in the data that are a result of some erroneous, unobservable, or incomplete process. Additionally, the observed range of the species may also be narrower than the actual range, and by using envelopes we are assuming that outside of the observed distribution there is an occurrence probability of zero. On the other hand, by using envelopes we are using the extremities of occurrence to define limits, which may result in broader classifications than using logisitic regression methods. Such reasons associate with data collection may help explain why there is a broad range of model performance, ranging from near perfect prediction, down to no better than random. The BIOCLIM method exemplifies some of this, as for the real species data it performed much worse than other methods, including our BBE method. The BIOCLIM method used all 19 predictors to classify the native range – and given so many variables, the distribution would need to be well-sampled to avoid overfitting. For the virtual species it worked reasonably well compared to the other methods, as it was able to fit a distribution based entirely on two variables included in the 19 available.

Another issue that challenges SDM application is how different backgrounds (extent and scale) select different variables to be included in the process. For example, when using a method like Maxent, the size of the background can change the environmental gradients being sampled, changing the shape of the response curves and predictor importance (and thus selection) (Merow et al., 2013). Background selection or delimitation is therefore a key component of model transferability when considering the effect on response curves, so often some form of ecological justification (e.g. accessible area) is required (Barve et al., 2011; Owens et al., 2013). Using biomes to select backgrounds seemed to perform well for the species used in this study, previously explored in Hill et al. (2017) when examining transferability between continents. The range-bagging technique largely avoids the issue of background selection, but future work should include to examine how sensitive performance of the GAMs (in the CHE and BBE methods) is to the background selection.

### Recommendations

While we have not performed an exhaustive investigation of all modelling parameters and decisions, our results suggest there are a few approaches that appear to be useful for predicting non-native distributions as part of evaluating risk posed by plant pests (and other species of concern). In particular, by being more “conservative” in the absence of knowledge of the most proximal variables, the ensemble methods of CHE and pairwise Maxent, and the range bagging approach, perform well. This finding also supports that when no standardised set of predictors is required, using iterative approaches to predictor selection can increase predictive performance Petitpierre et al. (2017). One advantage to using range bagging is that is does not require a background to be specified, which may be useful when there is limited or sparsely distribution occurrence data are available. Range bagging is also less computationally expensive than performing multiple GAMs to rank predictor pairs in the CHE method, but given an appropriate background, CHE performs well.

It still remains to be investigated how choices on thresholds for omitting variable pairs influences results, as well as incorporating other variables outside of climate – and how they are incorporated as pairs or triples etc. For all approaches, we advocate for use of as much distribution data as possible, especially when the species may be present in multiple geographical regions, to define the environmental space more completely (Broennimann and Guisan, 2008; Hill et al., 2017). Of course data should always be examined for environmental outliers that would particularly alter results using convex hulls or other envelope approaches. An important caveat here is that for species where models projected badly, they tended to do so regardless of the modelling approach. In these cases it is hard to justify the use of any correlative or envelope methods to project them. Further, as we have focused on non-native insect species, it is now important to examine more datasets to see if patterns of performance hold across different taxa, such as other animal pests, weeds and perhaps pathogens.

The goals of the risk mapping exercise are likely to define what trade-off of computational time and method exploration are appropriate (Breiner et al., 2018). For risk maps is in aiding industry preparedness, there is typically a preference of predictive accuracy. Therefore, increased model transferability at the expense of model interpretability is perhaps more important for effective rapid predictions (Merow et al., 2014), especially in the management of non-native species and biological invasions. This is especially important if the costs associated with overprediction (arising from more conservative models) translates to significantly less than the cost of the species invading (Jiménez-Valverde et al., 2011).

## Acknowledgements

Daniel Heersink wrote earlier versions of some of the code used in this study. Trevor Booth and Scott Foster provided valuable comments for this manuscript. Funding was provided by Centre for Excellence in Biosecurity Risk Analysis.

## Data and Code accessibility

The species distribution data, functions to fit CHE, BBE and the modified range bagging models, and a short tutorial are available as an R package, currently hosted at: https://github.com/mattecologist/che

